# *Nix* confers heritable sex-conversion in *Aedes aegypti* and *myo-sex* is needed for male flight

**DOI:** 10.1101/595371

**Authors:** Azadeh Aryan, Michelle Anderson, James K. Biedler, Yumin Qi, Justin M. Overcash, Anastasia N. Naumenko, Maria V. Sharakhova, Chunhong Mao, Zach N. Adelman, Zhijian Tu

## Abstract

A dominant and hemizygous male-determining locus (M locus) establishes the male sex (M/m) in the yellow fever mosquito, *Aedes aegypti*. *Nix* is a male-determining factor (M factor) in the M locus and its transient expression in females (m/m) results in partial masculinization. Here, we show that the *Nix* transgene alone was sufficient to convert females into fertile males, which continued to produce sex-converted progeny in subsequent generations. However, assisted mating with wild-type females was necessary, as the converted m/m males could not fly. Knockout of *myo-sex*, a myosin heavy chain gene and the only other protein-coding gene reported in the M locus, rendered wild-type males flightless. Thus, *Nix* alone converts female *Ae. aegypti* to fertile males and *myo-sex* is required for male flight. Only female *Ae. aegypti* mosquitoes bite and transmit disease-causing viruses. *Nix*-mediated female-to-male conversion is 100% penetrant and stable over many generations, indicating great potential for mosquito control.

---

Highly diverse primary signals serve as the master switches to initiate sex determination in insects (reviewed in Bachtrog *et al.*, 2014, Biedler and Tu, 2016). In some species, the initiating signals are female-determining factors that trigger female development. For example, a double dose of the X-linked signal elements in the fruit fly *Drosophila melanogaster* instigates female development in XX embryos (Salz and Erickson, 2010); in honeybees the heterozygosity of the *complementary sex determiner* (*csd*) gene initiates female development in diploid embryos produced by fertilized queen bees (Hasselmann *et al.*, 2008); and a W chromosome-linked piRNA gene determines female sex in ZW silkworms (Kiuchi *et al.*, 2014). In contrast, a dominant male-determining factor (M factor) serves as the primary signal that triggers male development in many other insects including mosquitoes and other non-Drosophila flies, beetles, and true bugs (Baker and Sakai, 1979, Hilfiker-Kleiner et al., 1994, Willhoeft and Franz, 1996, Shukla and Palli, 2014, Bachtrog *et al.*, 2014, Charlesworth and Mank, 2010). The M factor is located either on a Y chromosome or within a sex locus named the M locus on a homomorphic sex-determining chromosome, both of which are repeat-rich and thus difficult to study. Facilitated by recent advances in bioinformatics and genetic technologies, the M factor has been discovered in four dipteran insects including *Nix* in the yellow fever and dengue fever mosquito *Aedes aegypti* (Hall et al., 2015), *gYG2/Yob* in the African malaria mosquito *Anopheles gambiae* (Hall et al., 2016, Krzywinska et al, 2016), *Guy1* in the Asian malaria mosquito *An. stephensi* (Criscione et al., 2016), and *Mdmd* in the housefly *Musca domestica* (Sharma et al., 2017). None of these M factors are homologous to each other, indicating frequent turnover of the initiating signals for sex determination. However, through a cascade of events, these highly divergent primary signals are eventually transduced as sex-specific isoforms of conserved transcription factors doublesex (DSX) and fruitless (FRU) that program sexual differentiation. Thus, diverse primary signals in different species regulate the alternative, sex-specific splicing of *dsx* and *fru* pre-mRNAs, leading to sex-specific DSX and FRU protein isoforms.

Sex conversion resulting from a loss-of-function mutant male or a gain-of-function female would strongly indicate that an M factor is the master switch for sex determination. Such evidence has been presented only in species for which sex chromosome dosage compensation is not necessary (Hall et al., 2015, Sharma et al., 2017). For example, we have shown that somatic knockout of *Nix* in male embryos resulted in feminized *Ae. aegypti* adults with developing ovaries. Female embryos injected with a plasmid that contains the *Nix* ORF driven by a strong constitutive promoter (polyUb) developed into partially masculinized adults (Hall et al., 2015). However, full phenotypic sex conversion was not observed presumably due to somatic mosaicism, the transient nature of *Nix* expression, and/or the use of a different promoter. Thus, it remains unclear whether *Nix* alone is sufficient for conferring complete male sexually dimorphic traits and fertility.

Initial efforts failed to produce transgenic lines when *Nix* was under the control of the polyUb promoter. Thus, we sought to make transgenic mosquitoes that stably express a *Nix* transgene under the control of its native promoter. We isolated a 2.5 kb region upstream of the *Nix* gene, which was used to express the *Nix* transgene in the donor plasmid (Figure 1A). In addition to the native *Nix* ORF (N1), we also designed a second donor plasmid that contains a Strep II tag (Schmidt and Skerra, 2007) at the N-terminus (N2) to facilitate future biochemical studies and to distinguish between the endogenous *Nix* and the *Nix* transgene (Figure 1A). Three transgenic lines were obtained (Table S1–3) using the donor plasmids shown in Figure 1A. In subsequent analyses we focused on two of these (one line derived from each of the two constructs), hereto referred to as N1 and N2. The transgene insertion sites were identified by inverse PCR (Figure S1, primers are shown in Table S4). Bioinformatic mapping of the sequences flanking the N2 transgene revealed an insertion on chromosome 2 (Figure 1B), while the N1 transgene was present near the telomere on the 1p arm of chromosome 1 (Figure 1B), distantly linked to the native M locus, as confirmed by chromosomal florescent *in situ* hybridization (FISH, Figure 1C). To determine whether the native *Nix* promoter fragment recapitulated *Nix* expression in the context of these new genomic locations, we determined the transcription profile of the N2 transgene using primers that readily distinguish the N2 *Nix* transcript from the endogenous *Nix* transcript (Figure S2, Table S4). Like the endogenous *Nix*, N2 *Nix* transcription was observed from the onset of embryonic development starting at 2-4 hrs after egg deposition throughout all developmental stages. The N2 *Nix* transcript was detected in transgenic m/m pupae and adults but not in wild-type male pupae or adults. Also as expected, we did not observe endogenous *Nix* transcripts in transgenic m/m pupae or adults. Thus, the 2.5 kb *Nix* promoter that we isolated appears to function similarly to the native *Nix* promoter in the M locus in its temporal expression pattern.

**Figure 1.**
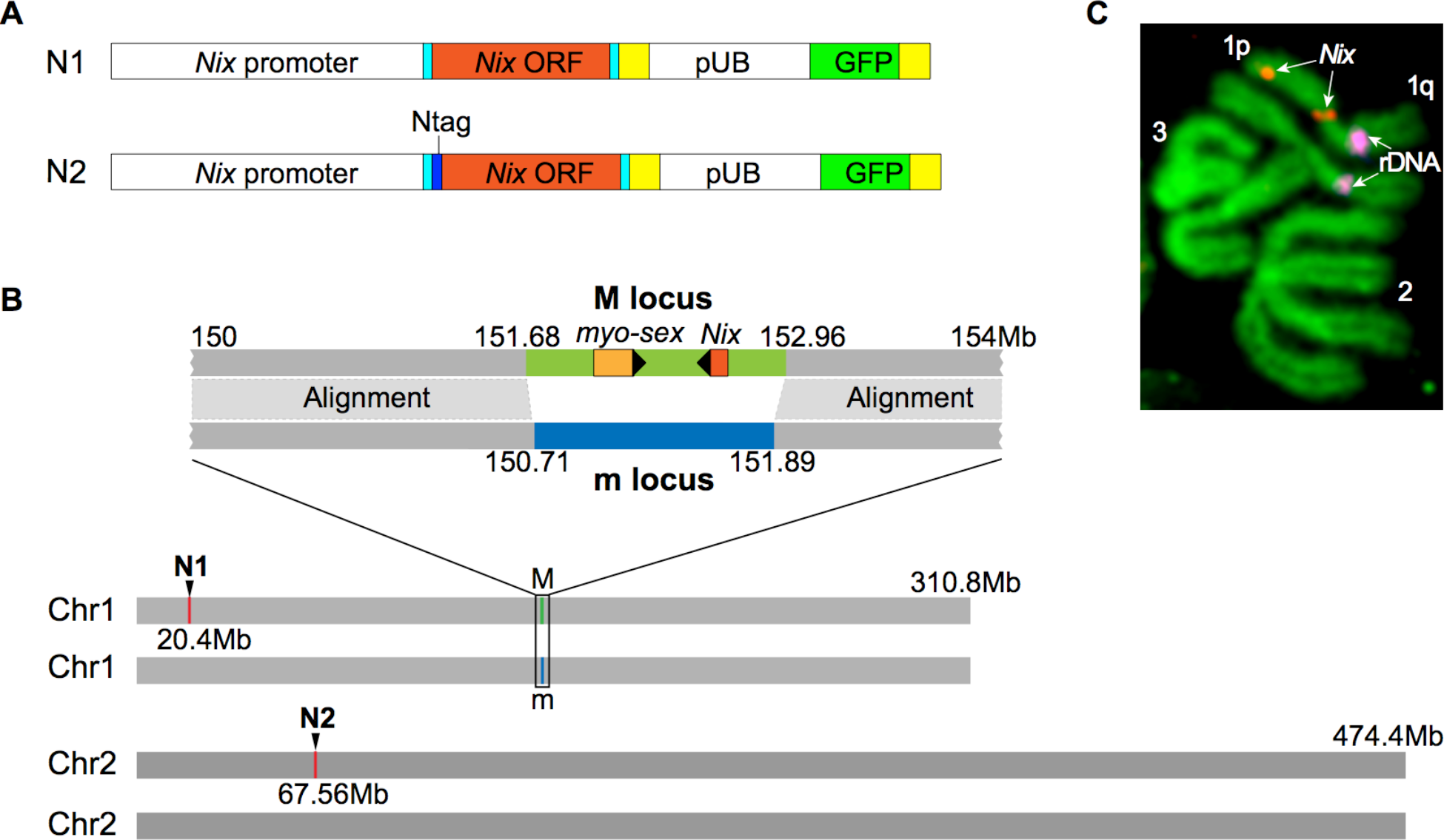
Transgenic lines that stably express the *Nix* transgene. **A)** Plasmid constructs used to create transgenic lines that stably express the *Nix* transgene. N1 was designed to express *Nix* from its own promoter. A 3.7 kb *Nix* sequence containing the approximately 2.5 kb promoter, the 5’ untranslated region (5’ UTR, light blue), the *Nix* ORF (orange), 3’ UTR (light blue) is followed by the SV40 polyadenylation signal (yellow). This *Nix* expression cassette is followed by the transformation marker cassette, GFP driven by the *Ae. aegypti polyubiquitin* promoter. These two cassettes are flanked by the Mos1 transposon arms, which are not shown (Coates et al., 1998). The N2 construct is identical to N1 except that a Strep-tag II (Schmidt and Skerra, 2007) was added to the N-terminus of the *Nix* ORF. **B)** A schematic drawing showing the insertion sites in the N1 and N2 transgenic lines, respectively. The relative position of M and m loci and the content of known genes in the M locus are also shown. **C)** Chromosomal *in situ* hybridization using the N1 plasmid as a probe showed a signal (red) on the p arm of chromosome 1 in addition to a signal in the known M locus (Matthews et al., 2018). The rDNA signal (magenta) was used as a landmark for the q arm of chromosome 1.

We next determined the phenotype of the genetic females (as indicated by the lack of the M locus including *myo-sex* and the native *Nix* gene, Figure S3) that contained the N1 or N2 *Nix* transgene. Four different genotypes are possible from crosses between transgenic males and wild-type females (Figure 2A). They can be distinguished by the presence or absence of the EGFP transgenic marker (N/+ vs +/+) and the presence or absence of the endogenous M locus (M/m vs m/m, Figure S3). All genetic females with the N2 transgene showed conversion to complete male sexually dimorphic features, including plumose antennae, external genitalia showing male gonocoxite and gonostylus, and internal sex organs such as testes and male accessory glands (Figure 2B-C, Figure S3). The same female-to-male conversion phenotype was also observed in the N1 line (Figure S4). Thus, the *Nix* transgene alone converted females into phenotypic males. The sex-converted males showed a slightly larger body size compared to the wild-type and transgenic M/m males, as indicated by their wing length measurement (Figure 2D, Figure S5, Table S5), which is used as a reliable proxy for body size (Gleiser et al., 2000). It is possible that subtle differences in *Nix* transgene expression and/or other factor(s) in the M locus may contribute to the wing length difference between sex-converted m/m males and M/m males. We performed RNAseq analysis of three biological replicates of pooled individuals of the four genotypes. As shown in Figure 2E, the overall transcription profile of the sex-converted m/m males was highly similar to wild-type M/m and transgenic M/m males, but clearly different from the wild-type m/m females. In addition, *dsx* and *fru* splicing was shifted towards the male isoforms, as confirmed by digital droplet RT-PCR (ddPCR, Figure S6).

**Figure 2.**
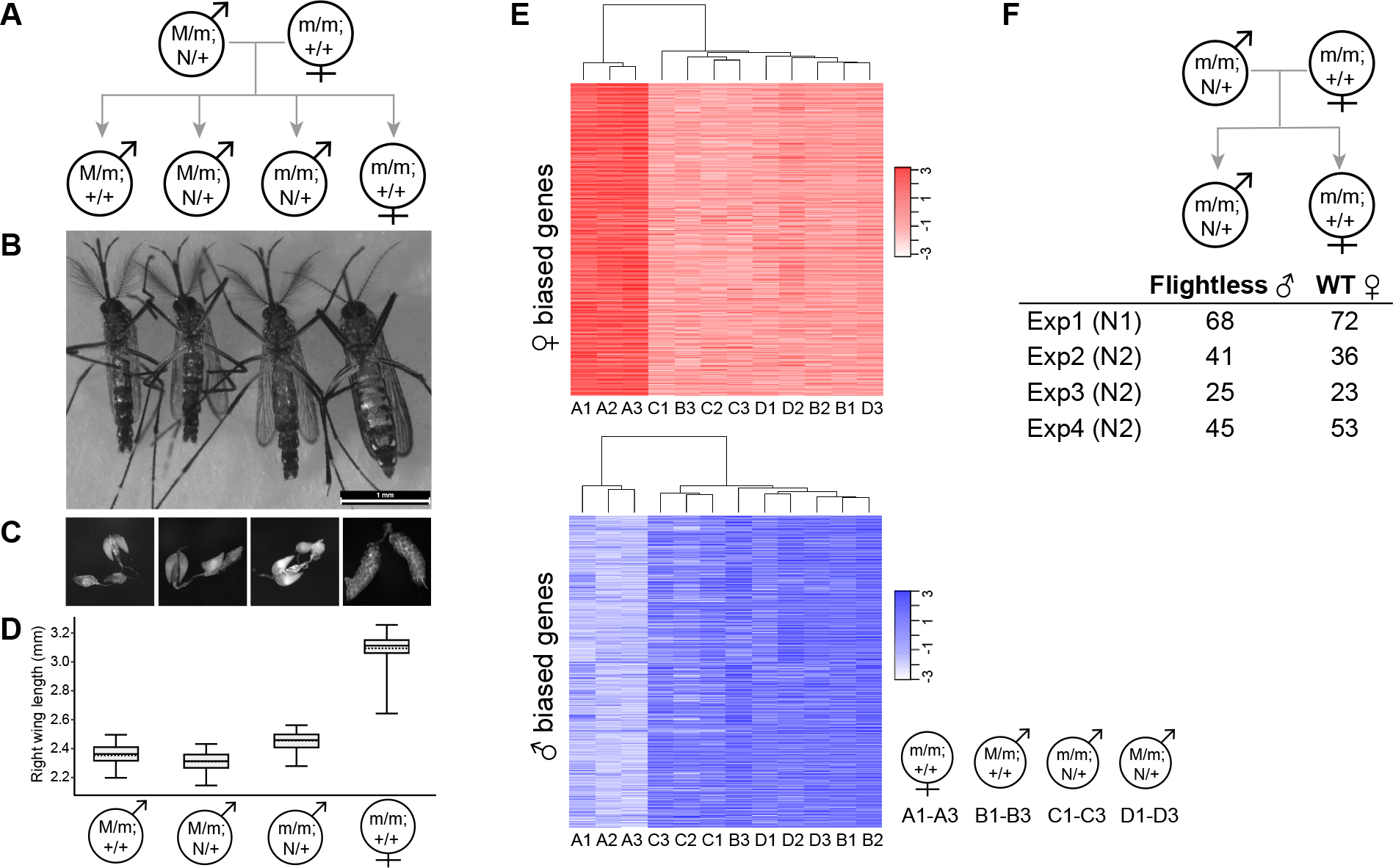
Nix transgene alone is sufficient to convert genetic females into fertile males with high penetrance and stability. **A)** Pedigree showing the four genotypes of progeny from a cross between transgenic (N/+) males (M/m) and wild-type (+/+, or non-transgenic) females (m/m). N denotes the N1 or N2 *Nix* transgene. In this particular mating scheme, N2 transgenic males were used and produced progeny shown in B-D. **B)** Representative individuals showing the phenotypes of the four genotypes shown in panel A. Genotyping was performed as shown in Figure S3. **C)** Reproductive organs from individuals of the four genotypes. Images were taken using the LAS V4.5 software suite with the zoom set to 1 (whole body) or 4 (dissected tissues). **D)** Length of the right wing of individuals of the four groups. Thirty individuals within each group were measured from the same cohort. Box plot, starting from bottom shows minimum values, 1^st^ quartile, median, 3^rd^ quartile, and maximum values using horizontal solid lines, with the mean indicated by a horizontal dashed line. As shown in Table S5, all pairwise comparisons were significant (p<0.0001), except for (M/m; +/+) vs. (M/m; N2/+) males (p=0.1982). **E)** RNAseq of biological triplicates of each of the four genotypes from a cross between N1 transgenic males and wild-type females. The log_2_(FPKM) expression level heatmap of female-biased (red) and male-biased (blue) genes are shown and clustering of the samples was based on the transcription profile. **F)** Assisted mating of the sex-converted flightless m/m males with wild-type females produces wild-type females and m/m flightless males in four independent experiments. Similar results were obtained for both N1 and N2 (indicated in brackets) transgenic m/m fathers. Genotype of the converted flightless males was confirmed (Figure S7).

Interestingly, all genetic females that were converted to phenotypic males due to the ectopic expression of the N1 or N2 transgene could not fly (Table 1, and a supplemental video at https://youtu.be/fwUqN5iKTi0). The majority of flightless males were not able to completely fold their wings; they could walk and sometimes jump but could never sustain flight. Wild-type males, N1 or N2 M/m males, and wild-type females displayed normal flight phenotypes. To assess the stability of these phenotypes, we screened for the four genotypes over multiple generations by crossing either (M/m, N1/+) or (M/m, N2/+) males with wild-type females. Between the two lines, we found 0 transgenic individuals that developed as females while scoring approximately 4300 transgenic males and 2000 flightless transgenic males (Table 1). As expected, the percentage of flightless males are higher in the N2 line than the N1 line because N1 is linked to the M locus while N2 is on a separate chromosome. However, the numbers of the four genotypes did not strictly follow the expected Mendelian segregation ratios in the N2 lines in some generations (Table S3). It is possible that some N2 male convertants may have died prior to adult emergence.

**Table 1.** Total number of progeny from crosses between transgenic males and wild-type females.

| Line | Transgenic ♂ | Non-transgenic ♂ | Transgenic ♀ | Non-transgenic ♀ | Transgenic flightless ♂ |
| --- | --- | --- | --- | --- | --- |
| N1 | 2696 | 829 | 0 | 2186 | 902 |
| N2 | 1610 | 1848 | 0 | 1771 | 1098 |

The latest screening was done for G<sub>13</sub> and G<sub>14</sub> for N1 and N2 lines, respectively. Detailed numbers from each generation are provided in Tables S2 and S3.

We then tested whether N1 or N2 m/m males were fertile, despite the absence of the M locus. As these converted males were flightless and flying is required during mating (Cator et al., 2011), they could not mate with females without assistance. Using assisted mating in four independent experiments that include both N1 and N2 lines, the sex-converted males produced viable progeny when crossed with wild-type females (Figure 2F). As there is no M locus at all in these sex-converted transgenic males (m/m; N/+), when mated with wild-type females (m/m; +/+), only two genotypes are possible in their progeny: (m/m; N/+) and (m/m; +/+), which will manifest as flightless sex-converted males and wild-type females, respectively. Indeed, flightless males and wild-type females were observed at an approximately 1:1 ratio in all four experiments, and genotyping results confirmed that these flightless males were indeed sex-converted genetic females (Figure S7). Thus, sex-converted phenotypic males are fertile and continued to produce sex-converted progeny, indicating that the *Nix* transgene alone is sufficient to convert females into fertile males and that this sex-conversion is heritable. These results, together with the fact that not a single transgenic individual developed a female phenotype over a combined 27 generations with thousands of individuals screened (Table 1), suggest that the female-to-male conversion conferred by the *Nix* transgene is highly penetrant and stable.

In addition to *Nix*, a myosin heavy chain gene, *myo-sex* (Hall et al., 2014) is also located in the M locus (Matthews et al., 2018). Thus far, *Nix* and *myo-sex* are the only protein-coding genes described in the M locus. Therefore, we hypothesized that the flightless phenotype observed in N1 (m/m) and N2 (m/m) males is caused by the lack of the *myo-sex* gene associated with the M locus. To test this hypothesis, we performed CRISPR/cas9-mediated knockout using sgRNAs designed to specifically target the hemizygous *myo-sex* gene (Figure 3). Following injection of pre-blastoderm embryos with a Cas9/sgRNA mixture, a significant portion of the surviving G_0_ males were flightless in three independent experiments. Despite the fact that only flying G_0_ males produced progeny when wild-type females were provided for mating, a significant portion of the G_1_ male offspring were also flightless in all three independent experiments (Figure 3). We sequenced plasmids cloned from PCR products from 21 flightless G_1_ males and identified mutations in the *myo-sex* gene in all of them. Thus, we conclude that the M locus gene *myo-sex* is required specifically for male flight.

**Figure 3.**
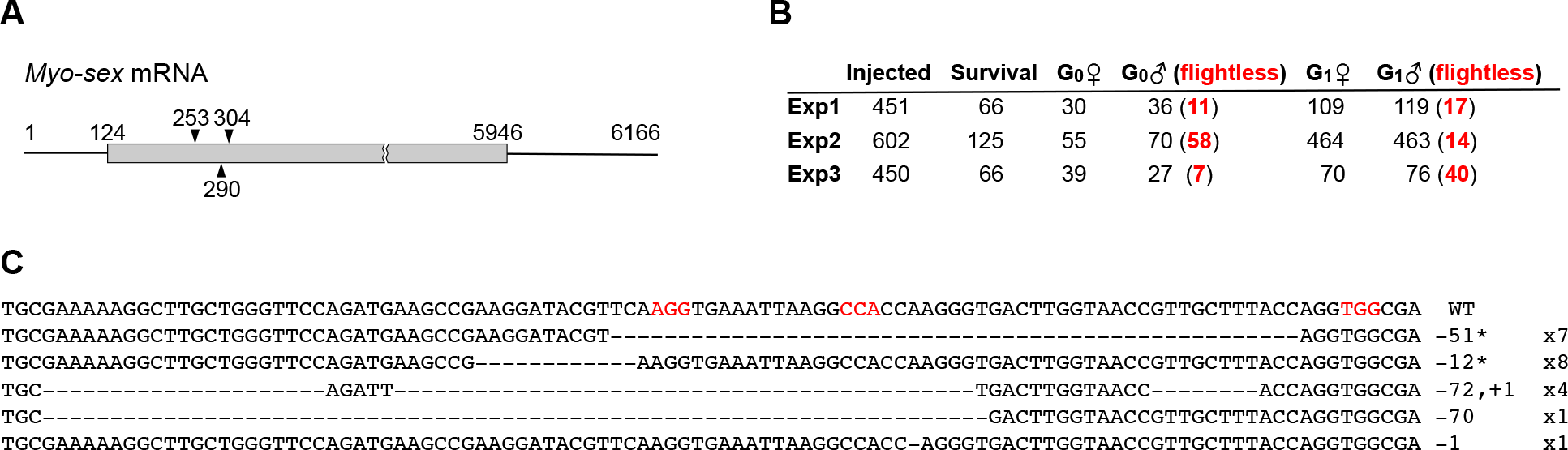
*Myo-sex* knockout results in flightless males. **A)** Three sgRNAs, starting at positions 253, 290, and 304, were used to target the *myo-sex* coding region, which spans positions 124 to 5946. **B)** CRISPR/Cas9-mediated *myo-sex* knockout produced flightless males in both the G_0_ and G_1_ generations in three independent experiments. G_1_ progeny was produced only from crosses between wild-type virgin females and sgRNA+Cas9-injected flying G_0_ males. **C)** Sequence analysis of mutations in CRISPR/cas9-edited flightless G_1_ progeny. Top line denotes the wild-type (WT) sequence; subsequent lines show various mutant sequences. The number of deleted (−) and inserted (+) bases and their occurrence are indicated to the right; in-frame mutations are indicated by an asterisk; deleted bases are denoted by dashes. The PAM sites for the three sgRNAs are highlighted in red.

Here we have shown that *Nix* alone fulfills the role of a dominant master switch for male determination in *Ae. aegypti*. The fact that *Nix* is transcribed at the onset of maternal-to-zygotic transition (Hall et al., 2015) and that *Nix* expression does not require any other factors in the M locus, as shown by the transcription of the *Nix* transgene in m/m females (Figure S2), indicate that *Nix* is the primary signal in the sex-determination pathway. Therefore, we conclude that *Nix* is indeed the M factor for sex determination in *Ae. aegypti*. In addition, we have shown that *myo-sex*, the only other protein-coding gene in the M locus, is needed for male flight. Although it remains unknown whether *Nix* and *myo-sex* together can transform genetic females into both fertile and flying males, this study defined the roles of the only two known protein-coding genes in the sex locus of a unique, homomorphic, sex-determining chromosome in an important insect species.

The high penetrance and stability we observed make *Nix* an attractive target for manipulation to introduce male bias to reduce the number of biting and egg-laying females. Although a full sex-conversion that produces fertile and flying males will potentially be more efficient in male production, converting genetic females into flightless males already successfully removes those females from the progeny. Coupling a *Nix* transgene with a conditional expression method (Fu et al., 2010), or a recently demonstrated gene drive strategy (Gantz et al., 2015; Kyrou et al., 2018) would enable the inheritance of the *Nix* transgene in almost all offspring, resulting in the removal or conversion of potentially all females. Such *Nix* transgenic lines that produce male-only progeny may be used as a population suppression method (Adelman and Tu, 2016) or to significantly reduce the cost and augment the scale of other mosquito control strategies that require the separation of males from females (Papathanos et al., 2018).

## Acknowledgements

We thank Kate Morton, Camden Delinger and Clare Morris for mosquito care and screening; Clemont Vinauger for advice on statistical analysis and producing one of the video clips; Karthikeyan Chandrasegaran for advice on wing length measurement; Brantley Hall and Giuseppe Saccone for comments; and Janet Webster and Jean Clarke for editorial suggestions. This work is supported by NIH grants AI123338, AI121853, and the Virginia Agriculture Experimental Station.

## Author contribution

ZT directed the study and wrote the paper with input and contributions from all authors. AA, MA, JB, YQ, CM, MS, and ZNA drafted relevant method sections and/or provided initial components of figures and/or critical revisions. JB and ZT isolated the *Nix* promoter and designed the *Nix* constructs. AA, JO, MA, and ZNA generated the *Nix* transgenic lines. JB performed inverse PCR and statistical analysis. JB and YQ performed RT-PCR and ddPCR. AA and MA performed forced mating and *myo-sex* knockout. YQ and MA performed PCR-based genotyping. AA and MA documented the phenotypes with the help of undergraduate assistants. MA and YQ prepared RNAseq libraries. CM performed RNAseq data analysis and drew Figures 1–3. AN and MS performed *in situ* hybridization. MA and AA videotaped the flightless males and CM edited the video.

## Supplemental Materials

### Materials and Methods

#### Identification of the *Nix* promoter region

To isolate the promoter/upstream sequence of *Nix*, PCR was performed using two primers (Table S4), one of which contained the *Nix* ORF and the other was from approximately 3 kb upstream. PCR was performed with an Arktik Thermal Cycler (Thermo Fisher Scientific) using *Ae. aegypti* (Liverpool) genomic DNA and the Q5 DNA Polymerase (NEB) according to the manufacturer’s protocol, generating a single amplicon of approximately 3 kb. PCR products were cloned using the PCR-Blunt II-TOPO Cloning Kit (Thermo Fisher Scientific) and JM109 Competent Cells (Promega). A 2575 bp sequence upstream of the ORF, a consensus derived from 6 clones, was used as the promoter. The ORF and the 5’ and 3’ UTRs of *Nix* were previously determined (Hall et al., 2015).

#### Transgenic constructs

As shown in Figure 1A, the N1 construct was designed to express *Nix* from its own promoter (Figure 1A). A 3.7 kb sequence containing the 2575 bp promoter described above (including the 102 bp 5’ UTR), the 864 bp *Nix* ORF, and the 19 bp *Nix* 3’ UTR, followed by the SV40 polyadenylation/termination signal sequence was synthesized (Epoch Life Science, Inc.), and cloned into the pM2_pUB_EGFP vector (Anderson et al., 2010) using the PstI/AsiSI restriction sites. The pM2_pUB_EGFP vector consists of Mos1 transposase substrate arms and the EGFP transformation marker driven by the *Ae. aegypti polyubiquitin* promoter (Anderson et al., 2010). The N2 construct was designed to allow *NIX* protein purification and identification via an N-terminal Strep-tag II (Trp-Ser-His-Pro-Gln_phe-Glu-Lys, Schmidt and Skerra, 2007) (Figure 1A). The N2 construct is identical to N1 except that the Strep-tag II was added to the N-terminus of the *Nix* ORF (synthesized by Epoch Life Science Inc.).

#### Mosquito rearing and Mos1-mediated transformation

*Ae. aegypti* mosquitoes (Liverpool strain) were maintained at 28°C and 60-70% humidity, with a 14/10 hour day/night light cycle. Adult mosquitoes were maintained on 10% sucrose and blood-fed using artificial membrane feeders and defibrinated sheep’s blood (Colorado Serum Company; Denver, CO). Donor plasmid was co-injected at 0.5 μg/μl with the Mos1 helper plasmid, pGL3-PUbMos1 at 0.3 μg/μl into one hour old embryos (Coates et al., 1998). Surviving G_0_ females were mated to Liverpool males in pools of 20-25. G_0_ males were mated individually to 5 Liverpool females and then merged into pools of 15-20 males. G_1_ larvae were screened for GFP fluorescence using a Leica M165 FC fluorescence microscope. Positive G_1_ individuals were out-crossed to Liverpool females to ensure that all transgene cassettes were stably inherited to the G_2_ generation. Images were taken of pupae and mature adults (7-10 days post emergence) using the LAS V4.5 software suite at a Zoom of 1.6 (adults, pupae) or 4 (heads, genitalia, dissected tissues) and the following settings: Gain 1, Gamma 1, greyscale (white light) or pseudocolor (509nm).

#### Inverse PCR for transgene insertion site determination

Inverse PCR was used to determine the insertion site of the transgenic cassette for the N1 (generation 10) and N2 (generation 7) transgenic mosquito lines. Genomic DNA was isolated from three male adult individuals for each line, using the Quick-DNA Miniprep Kit (Zymo Research) according to the manufacturer’s protocol, and with an elution volume of 50 ul H_2_O. Three restrictions enzymes (Thermo Fisher), HpaI, MspI, and Bsp143I, were used to digest genomic DNA from each individual of each transgenic line. Approximately 1 ug DNA was digested for 8 hrs using 4 ul enzyme in a 60 ul reaction volume. Digested DNA was purified using the illustra GFX PCR DNA and Gel Band Purification Kit (GE Helathcare) with an elution volume of 50 ul H_2_O. Approximately 500 ng DNA was ligated in a 400 ul reaction volume with 1 ul T4 DNA Ligase (3 Weiss Units) (Promega) overnight at 16° C. DNA was purified using the illustra GFX PCR DNA and Gel Band Purification Kit (GE Healthcare) with an elution volume of 30 ul H_2_O. PCR was performed using 1 ul of the purified DNA in a 50 ul reaction volume with Q5 DNA Polymerase (NEB), in a T100 Thermal Cycler (Bio-Rad), with primers (Table S4) specific for the Mos1 Transposable Element Right Hand Arm, which is located at the 5’ terminus of the transgenic cassette (see Figure 1A for transgenic constructs). One specific PCR product was observed for each of the 3 enzymes used, for both lines. Sequencing results for all PCR products were the same for all individuals within each line, consistent with a single insertion site for each of the transgenic lines.

#### Chromosome fluorescent *in situ* hybridization

Slides of mitotic chromosomes were prepared from imaginal discs of 4^th^ instar larvae from the N1 transgenic line following published protocols (Timoshevskiy et al., 2012). Fluorescent *in situ* hybridization was performed using the N1 plasmid construct (Figure 1A) as a probe and 18S rDNA as a landmark for the q arm of chromosome 1. The N1 plasmid probe was labeled by nick translation (Invitrogen Corporation, Carlsbad, CA, USA), with Cy3-deoxyuridine 3-triphosphate (dUTP) (Enzo Life Sciences Ltd Farmingdale, NY, USA). 18S rDNA was labeled by PCR reaction (Bioline, Taunton, MA, USA) with Cy5-dUTP (Enzo Life Sciences Ltd). Chromosomes were counterstained with Oxasole Yellow (YOYO-1) iodide and mounted in Prolong Gold Antifade (Invitrogen Corporation, Carlsbad, CA, USA). Slides were analyzed using a Zeiss LSM 510 Laser Scanning Microscope (Carl Zeiss Microimaging, Inc., Thornwood, NY, USA) at 1000x magnification.

#### Endogenous and transgenic *Nix* transcription profile

Transgenic *Nix* GFP positive males from line N2 (M/m; N2/+) were crossed with Liverpool strain females and blood fed. To collect aged embryos ~20-30 females were placed into 50 ml conical tubes with a wet cotton ball and a disk of filter paper at the bottom. Females were allowed to lay over the interval time and then removed from the tubes and eggs allowed to mature to the desired age. Embryos were transferred from the filter paper to a 1.5 ml tube with a fine paint brush and snap frozen in liquid nitrogen at the appropriate time. Embryos (n≥100) were collected for the following time points: 0-1 hr, 2-4 hr, 4-8 hr, 8-12 hr, 12-24 hr, and 24-36 hr. More eggs were collected from the same cage and hatched in order to collect siblings at all four larval stages (n=50). One day and two day old pupae as well as 2-4 day post emergence adults were snap frozen individually in 1.5 ml tubes so they could be genotyped by PCR before further processing (Figure S3). RNA was isolated using Quick-RNA Miniprep (Zymo Research, Irvine, CA). cDNA was then synthesized using the SuperScript RT kit (Life Technologies, Carlsbad, CA). RT-PCR was performed using Phire II DNA polymerase (Thermo Fisher Scientific, Waltham Massachusetts) and primers are listed in Table S4. We took advantage of the Strep-tag II inserted between the 5’UTR and the ORF of the *Nix* in the N2 construct to design primers that only amplify cDNA from the endogenous *Nix* transcript (Table S4). Similarly, primers were also designed to only amplify the N2 transgenic *Nix* transcript (Table S4).

#### Wing length measurement and statistical analysis

Wing length values were measured for 30 individuals from line N2, eclosed from the same cohort of G14 larvae. Right wings were detached from one day post emergence adults, and mounted on a slide. A photograph was taken of each wing using a Leica DFC3000 G camera mounted on a Leica M165 FC Fluorescent Stereo Microscope. A 2 mm scale bar was photographed to standardize size measurements. ImageJ was used to measure the wing length as the distance from the anal lobe to the wing tip (Gleiser et al., 2000; Van Handel and Day, 1989; Figure S5). Wing length values for each phenotype are reported in Box Plots (Figure 2D) and shown in Table S5. One-way Analysis of Variance was performed using all samples, resulting in a P value of <0.0001. The Tukey Simultaneous Test for Difference of Means was performed resulting in adjusted P values <0.0001 for all pairwise comparisons except for transgenic males vs wild-type males (p = 0.1982). A Test for Equal Variances was performed using both Levene’s method and the multiple comparisons method.

#### RNASeq

Two to four day post-emergence adult siblings from the N1 transgenic line were snap frozen in liquid nitrogen and preserved at −80°C until the time of extraction. RNA was extracted using the Quick-RNA Miniprep kit (Zymo Research, Irvine, CA). Triplicate RNASeq libraries were prepared for each of the four genotypes (Figure 2E) using the NEBNext Ultra RNA Library Prep Kit for Illumina with the NEBNext Poly(A) mRNA Magnetic Isolation Module (New England Biolabs; Ipswich, MA) and multiplexed into 1 lane of a HiSeq 2500.

#### RNAseq data analysis

RNAseq reads from the wild type and transgenic mosquito samples were aligned using Tophat2 (v2.1.1) to the *Ae. aegypti* L3 reference genome. The resulting BAM files were sorted and indexed. MarkDuplicates from the Picard tool kit version 1.119 was used to identify and remove PCR duplicates (http://broadinstitute.github.io/picard/). Cufflinks version 2.2.1 was used to assemble transcripts and estimate the relative abundances of the transcripts (Robert et al., 2011). Transcription levels were estimated as Fragments Per Kilobase per Million mapped reads (FPKM). The reference transcript file (AaegL3.3.gtf) was downloaded from VectorBase (vectorbase.org). Cuffdiff, a component of Cufflinks was used to normalize and compare the transcript expression levels between samples (Trapnell et al., 2013). The log_2_(FPKM) expression level heatmaps (Figure 2E) were generated using the heatmap.2 function in R’s gplots package (https://www.rdocumentation.org/packages/gplots/). Both columns (samples) and rows (genes) were clustered using the default hierarchical clustering settings. Row scaling was applied and the row Z score values were used for the color scale.

#### Droplet digital PCR for *doublesex* and *fruitless* isoforms and data analysis

Total RNA was isolated from each of the four genotypes using the ZymoResearch Quick-RNA MiniPrep Kit (Irvine, CA) according to the manufacturer’s protocol. Three adult individuals (biological replicates) were used for each sample and all were siblings from generation 9 of the N1 transgenic line. cDNA was synthesized in a 20 ul reaction volume with approximately 500 ng total RNA and random hexamers, using the Invitrogen SuperScript III First-Strand Synthesis Super Mix (Waltham, MA) according to the manufacturer’s protocol, and the 20 ul completed cDNA reaction was diluted to a total of 60 ul with H_2_O. cDNA quality was checked by PCR using primers specific for ribosomal protein S7 (Rps7). These primers span a 110 nt intron which should yield a 501 or 611 bp PCR product for cDNA or a genomic DNA template, respectively. Primers were ordered from Sigma Aldrich (St Louis, MO). *Ae. aegypti* (*Liverpool*) male genomic DNA and H_2_0 was used as templates for positive and negative controls, respectively. PCR was performed using Phire HS II DNA polymerase from Thermo Fisher Scientific (Waltham, MA) according to the manufacturer’s protocol. One microliter of the cDNA reaction was used in a total of 20 ul PCR reaction volume and run for 30 cycles on a Bio-Rad MyCycler thermal cycler (Hercules, CA). Products were size-separated on a 1% agarose gel by electrophoresis, using a 100 bp step ladder from Promega (Madison, WI). Droplet digital PCR (ddPCR) was performed using 1 ul of the cDNA reaction with the Bio-Rad QX100 ddPCR machine (Hercules, CA) according the manufacturers protocols. Taqman assays were designed to detect sex-specific *doublesex* and *fruitless* isoforms (Table S4). The gene AAEL002401 was used as an internal reference for gene expression (Hall et al., 2015; Hu and Tu, 2018). Probes were ordered from Biosearch Technologies (Petaluma, CA) and primers were ordered from Sigma Aldrich (St Louis, MO). Expression values are reported as the mean +/− the standard error of the mean (Figure S6). One-way Analysis of Variance was performed using all samples for each assay and the Tukey Simultaneous Test for Difference of Means was performed for all pairwise comparisons (Figure S6). A Test for Equal Variances was performed using both Levene’s method and the multiple comparisons method.

#### Forced mating of flightless males

One week old virgin Liverpool strain females were blood fed to engorgement and incubated at 4 °C for 3-4 hours. Flightless males were added to the cage of cold-anesthetized females and allowed to mate as the females slowly woke up, over a period of about 5 minutes. Eggs were collected two days later and hatched upon maturation.

#### CRISPR/Cas9 mediated *myo-sex* deletion

Genomic regions of the *myo-sex* gene were manually searched for the presence of NGG (PAM), where N is any nucleotide. Selected target sites are 5’-GAAGCCGAAGGATACGTTCA<u>AGG</u>-3’, 5’-GTAACCGTTGCTTTACCAGG<u>TGG</u>-3’, and 5’-GGTTACCAAGTCACCCTTGG<u>TGG</u>-3’, with PAM sites underlined. sgRNAs were generated as previously described (Bassett et al., 2013) using primers listed in Table S4, *in vitro* transcribed using MEGAscript T7 kit and purified using the MEGAclear kit (Thermo Fisher Scientific; Waltham, MA). RNAs were aliquoted and stored at −80°C. Cas9 mRNA was *in vitro* transcribed as previously described (Basu et al., 2015) and used for the first two knockout experiments (Figure 3B). For the third replicate, Cas9 (ARCA) mRNA (Trilink Biotechnologies; San Diego, CA) was used. Three rounds of injections were performed with Cas9 mRNA at 0.6μg/μL and all three sgRNAs at 0.1 μg/μL each for ~500 embryos per replicate experiment. G_0_ males displayed a noticeable phenotype and were grouped by ability to fly, then crossed with Liverpool females. Mutations in the flightless male G_1_ were verified using the Phire II Animal Tissue Direct PCR kit (Thermo-Fisher Scientific; Waltham, MA), following the dilution procedure utilizing a single leg for the DNA template. Primers used for mutation screening are listed in Table S4. PCR amplicons were purified using the NucleoSpin Gel and PCR Clean-up kit (Macherey-Nagel; Düren, Germany) and sequenced.

---

**Table S1.** Plasmid injection and screening of G1 for transformants

| Donor plasmid | Mos1 helper | # embryos injected | # G0 (%survivors) | # pools | # of G1 screened | Pools with GFP+ G1 progeny (#) |
| --- | --- | --- | --- | --- | --- | --- |
| pMos-*Nix*-PUBGFP (N1) | pGL3-PUBMos1 | 1053 | 130 (12.3%) | 7 | ~13180 | P4<br>(5 normal male) |
| pMos-*Nix*-PUBGFP (N1) | pGL3-PUBMos1 | 1107 | 62 (5.6%) | 4 | ~7970 | N3<br>(12 normal male) |
| pMos-Ntag*Nix*-PUBGFP (N2) | pGL3-PUBMos1 | 1084 | 110 (10.1%) | 5 | ~10600 | P4<br>(4 flightless♂, 1 normal ♂) |

**Table S2.**
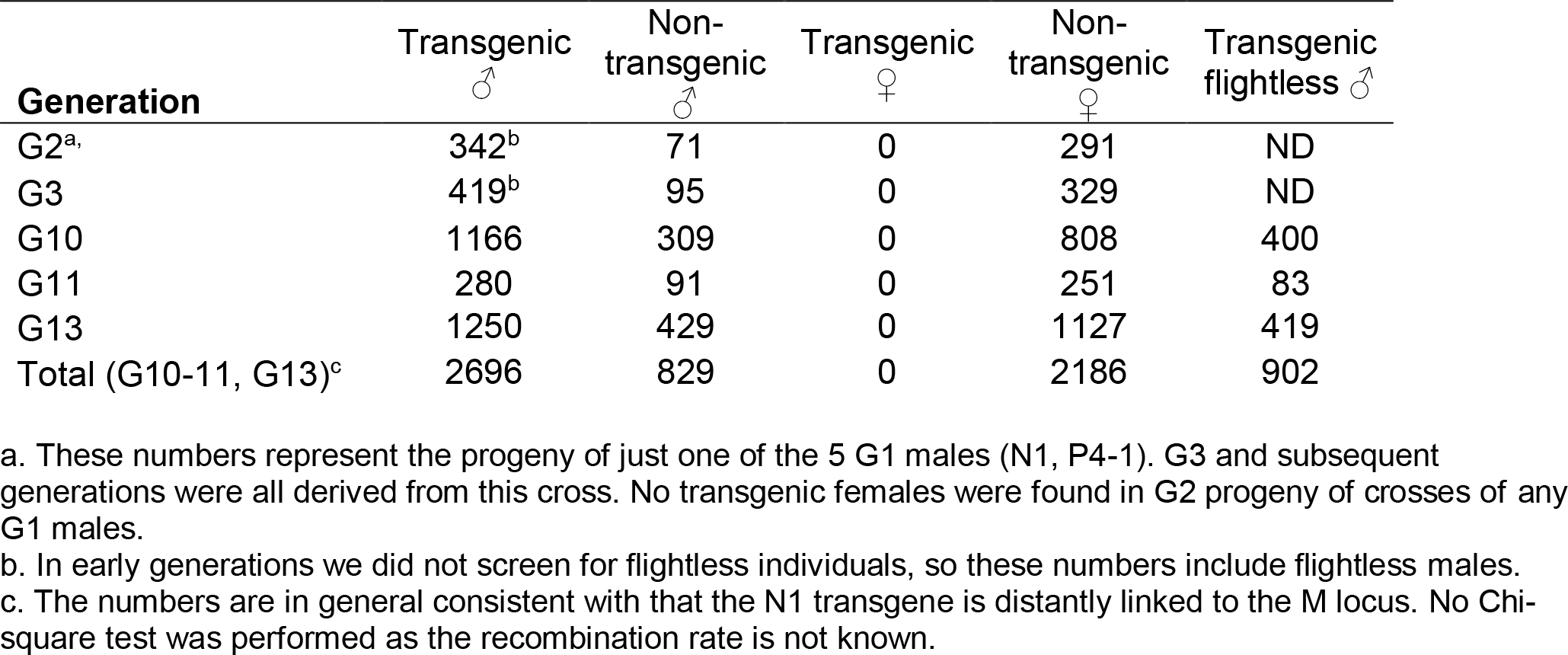
Screening of the N1 transgenic line

| Generation | Transgenic ♂ | Non-transgenic ♂ | Transgenic ♀ | Non-transgenic ♀ | Transgenic flightless ♂ |
| --- | --- | --- | --- | --- | --- |
| G2<sup>a</sup>, | 342<sup>b</sup> | 71 | 0 | 291 | ND |
| G3 | 419<sup>b</sup> | 95 | 0 | 329 | ND |
| G10 | 1166 | 309 | 0 | 808 | 400 |
| G11 | 280 | 91 | 0 | 251 | 83 |
| G13 | 1250 | 429 | 0 | 1127 | 419 |
| Total (G10-11, G13)<sup>c</sup> | 2696 | 829 | 0 | 2186 | 902 |

a. These numbers represent the progeny of just one of the 5 G1 males (N1, P4-1). G3 and subsequent generations were all derived from this cross. No transgenic females were found in G2 progeny of crosses of any G1 males.
b. In early generations we did not screen for flightless individuals, so these numbers include flightless males.
c. The numbers are in general consistent with that the N1 transgene is distantly linked to the M locus. No Chi-square test was performed as the recombination rate is not known.

**Table S3.** Screening of the N2 transgenic line

| Generation | Transgenic ♂ | Non-transgenic ♂ | Transgenic ♀ | Non-transgenic ♀ | Transgenic flightless ♂ |
| --- | --- | --- | --- | --- | --- |
| G2<sup>a</sup> | 242<sup>b</sup> | 152 | 0 | 124 | ND |
| G3 | 142 | 168 | 0 | 151 | 117 |
| G7 | 786 | 749 | 0 | 763 | 348 |
| G8 | 138 | 188 | 0 | 154 | 122 |
| G9<sup>c</sup> | 342 | 531 | 0 | 501 | 301 |
| G14 | 202 | 212 | 0 | 202 | 210 |
| Total (G3, G7-G9, G14)<sup>d</sup> | 1610 | 1848 | 0 | 1771 | 1098 |

a. These numbers represent the progeny of the only flying G1 males (N2, P4-2).
b. In this generation we did not screen for flightless individuals, so these numbers include flightless transgenic males.
c. 37 dead males were not screened.
d. As N2 is inserted in chromosome 2, we expected a 1:1:1:1 ratio among the four possible genotypes, which was observed in G14 (Chi-square test, $p=0.94$ ). However, this expectation was not met in earlier generations (Chi-square test, $p<0.05$ ).

**TableS4.**
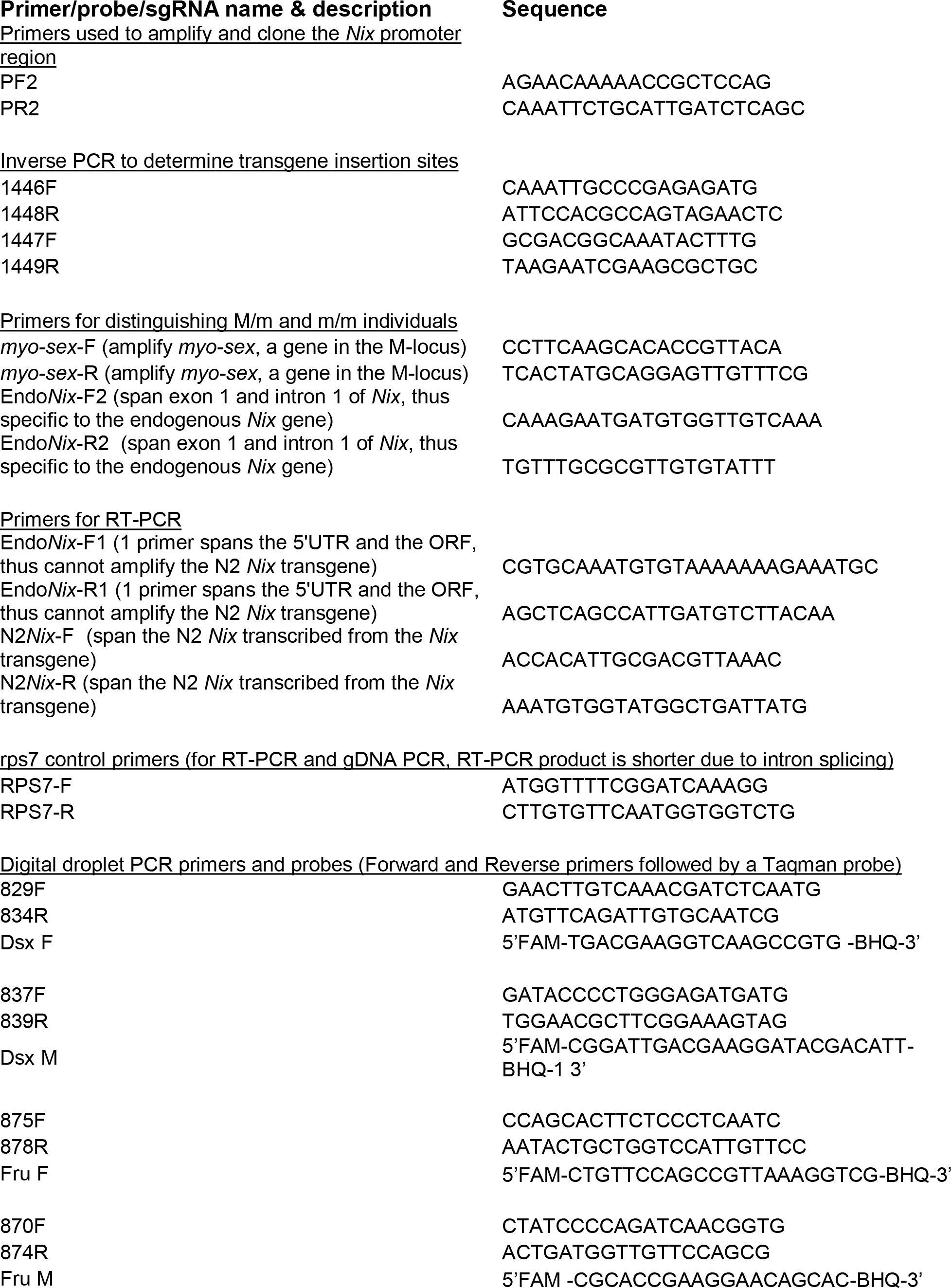
Primers, probes, sgRNAs used in this study

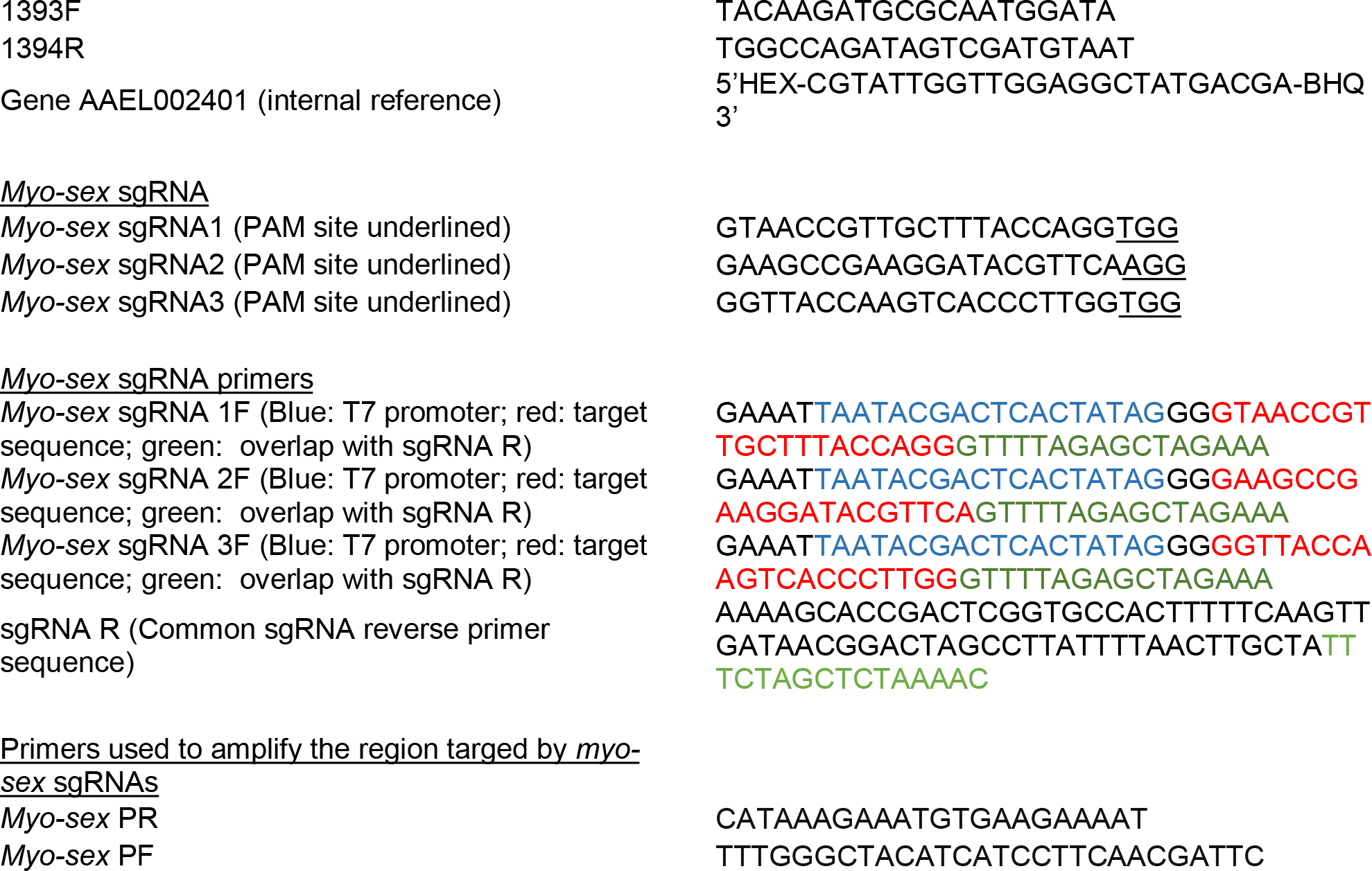

**Table S5.**
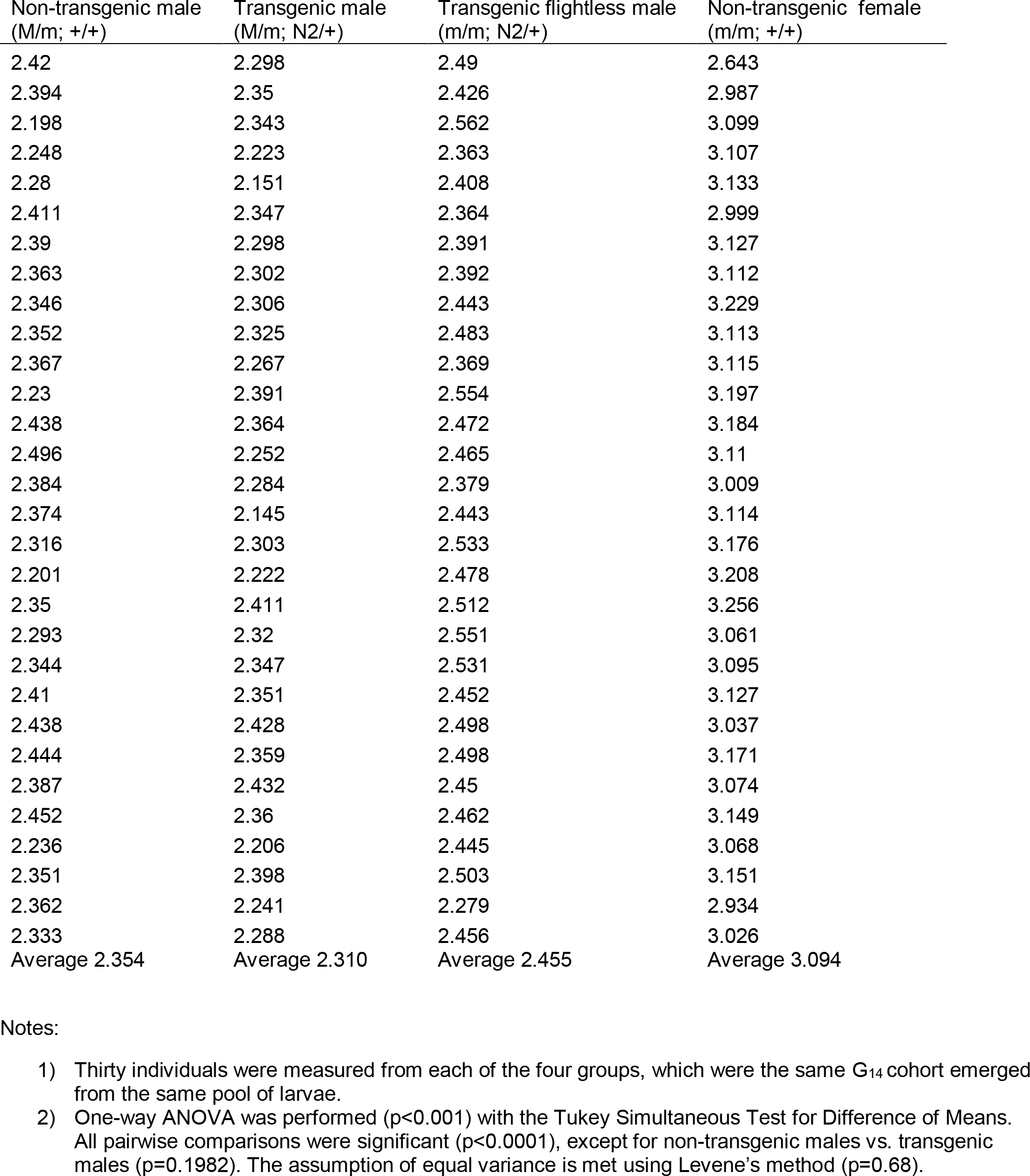
Length (millimeter) of the right wing of individuals of the N2 transgenic line

**Figure S1.**
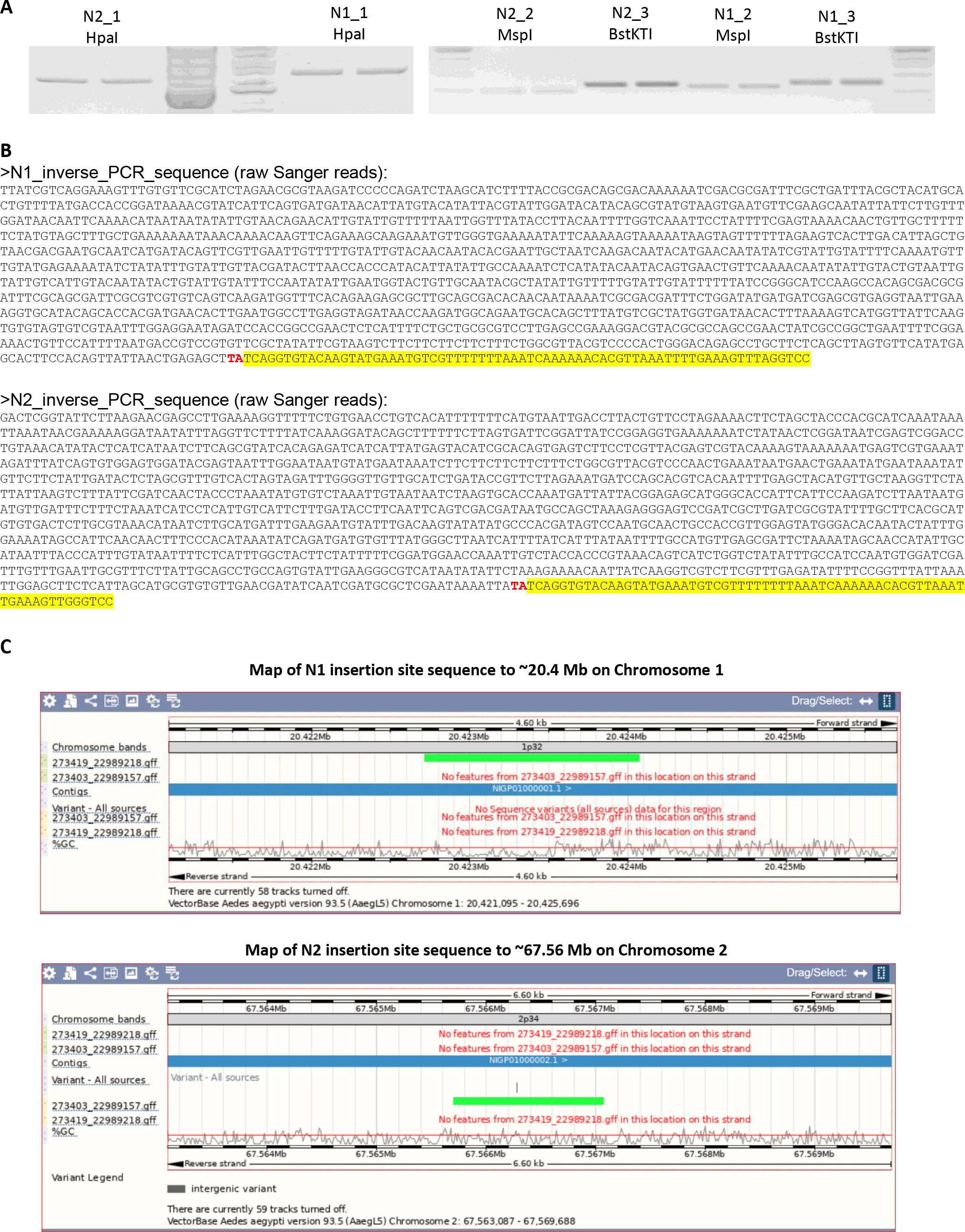
Inverse PCR to identify transgenic cassette genomic insertion sites. **A)** 1% agarose gel image showing PCR products resulting from PCR on ligated genomic DNA isolated from transgenic individuals. Genomic DNA of three separate individuals (_1, _2, and _3) from each transgenic line (N1 and N2) were digested with 3 restriction enzymes HpaI, MspI, and BstKTI. **B)** Genomic insertion site sequence identified by cloned and sequenced PCR products. For each transgenic line, all 3 reactions with different restriction enzymes resulted in identical sequences for the same genomic loci. “TA” target site (red highlight), and 5’ end of transgenic cassette, including the Mos1 Right Arm (Yellow highlight), are indicated. **C)** Mapping of transgenic insertion sites to the AaegL5 Genome (Vectorbase.org) by BLAST of sequences of cloned inverse PCR products.

**Figure S2.**
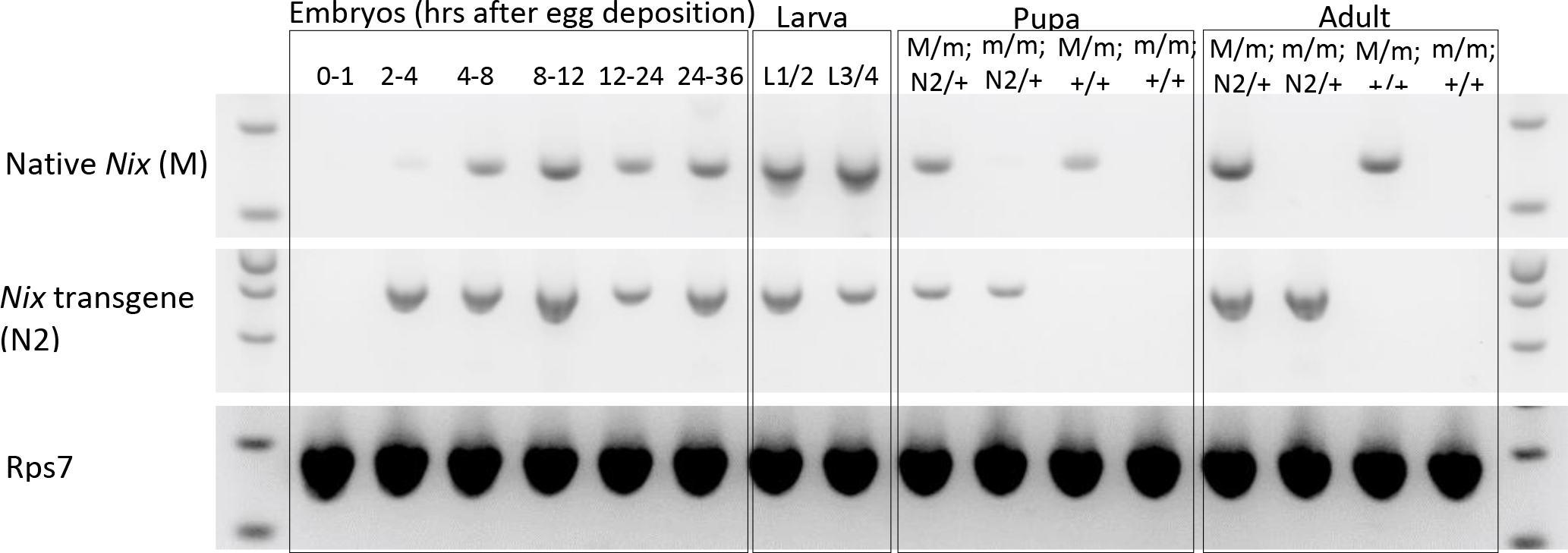
RT-PCR showing the transcription profile of the N2 transgene and the endogenous *Nix* gene. L1/2 indicates mixed sex first and second instar larva. L3/4 indicates mixed sex third and fourth instar larvae. Genotypes of the pupa and adults was determined by the presence and absence of the EGFP transformation marker and by the presence and absence of endogenous *Nix* and *myo-sex* (Figure S3D).

**Figure S3.**
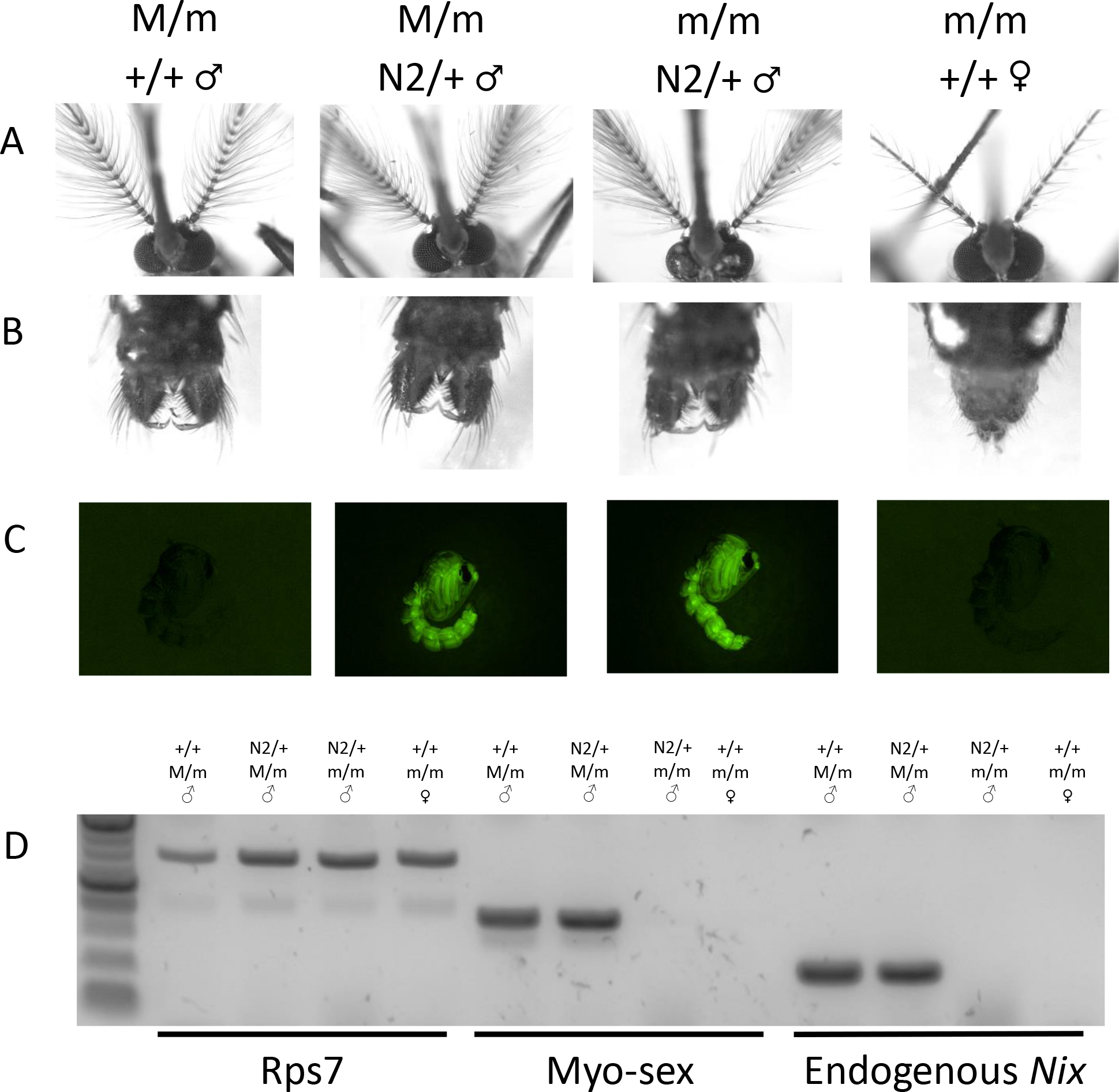
Images of sexually dimorphic features of all four genotypes and methods used to determine the genotypes. White light images of heads (A) and genitalia (B) of the four phenotypes observed in N2 transgenic *Nix* lines. ♂/♀reflects the observed phenotype. Images taken using the LAS V4.5 software suite with zoom set to 4 and the following settings: Gain 1, Gamma 1, greyscale (white light) or pseudocolor (509nm). C). The presence of a *Nix* transgene (N2) is determined by the presence EGFP marker. +/+ indicate wild-type or the lack of a transgene. D). The absence of the M locus, as indicated by the absence of two M locus genes *myo-sex* and the endogenous *Nix*, is used to indicate the m/m genotype.

**Figure S4.**
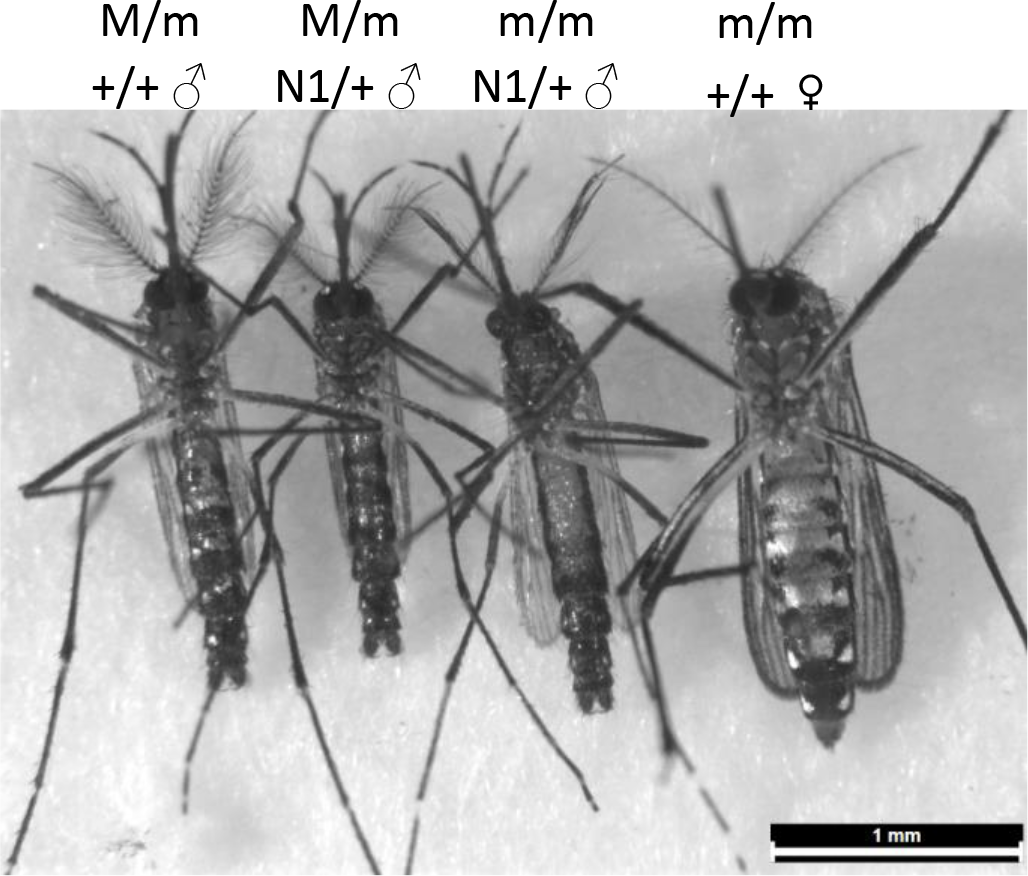
Full body images of the four phenotypes observed in N1 *Nix* transgenic lines. Representative individuals of each phenotype for the N1 line. Images were taken using the LAS V4.5 software suite with zoom set to 1, and the following settings: Gain 1, Gamma 1, greyscale

**Figure S5.**
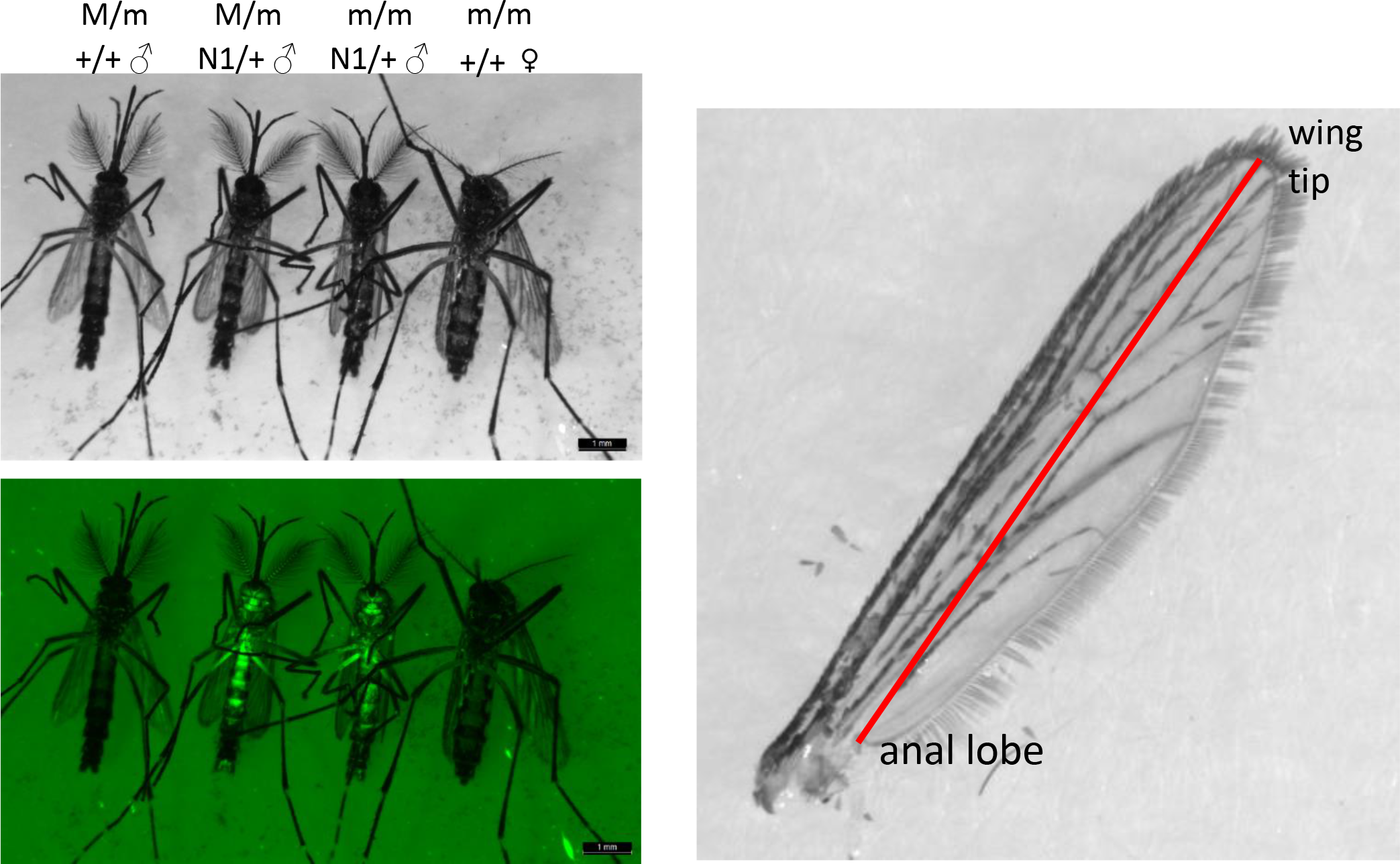
Method to measure the wing length of adult mosquitoes. Top left panel is the black and white image of the four genotypes to be measured. Bottom left panel is the florescent image of the same individuals. The right panel illustrates that the measurement is taken from the anal lobe to the wing tip.

**Figure S6.**
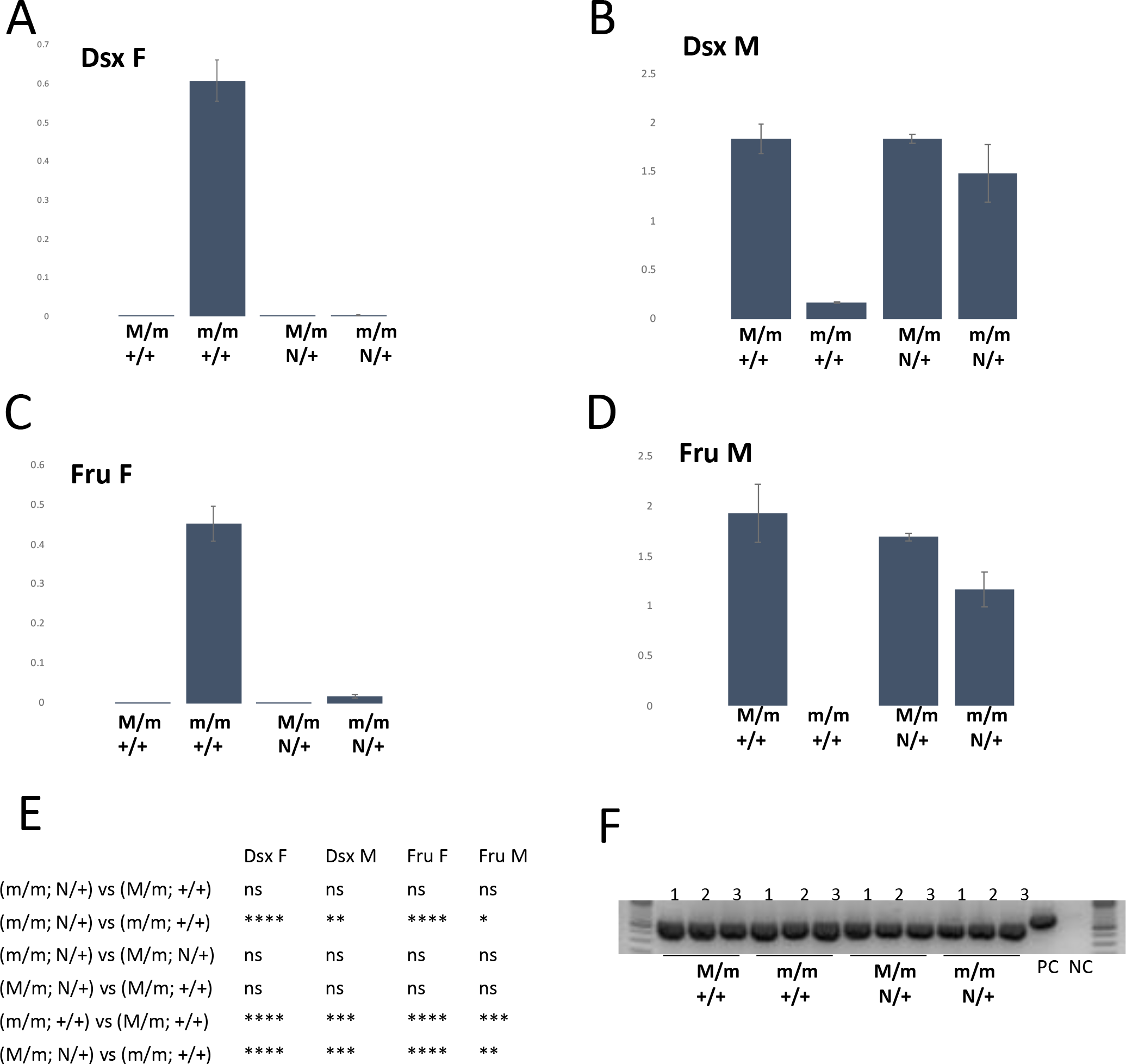
Digital droplet PCR assays for the N1 transgenic line show *doublesex* and *fruitless* isoform levels in sex-converted flightless males (m/m; N/+) are similar to wild-type males (M/m; +/+). **A)** *Doublesex* female isoform **B)** *Doublesex* male isoform. **C)** *Fruitless* female isoform **D)** *Fruitless* male isoform. Values shown are the mean +/− SEM. The gene AAEL002401 was used as an internal reference for gene expression (Hall et al., 2015; Hu and Tu, 2018). **E)** Summary of statistical significance from all pairwise comparisons using a one=way ANOVA followed by Tukey Simultaneous Tests of Differences of Means (p > 0.05, ns; p </= 0.05, *; p </= 0.01, **; p </= 0.001, ***; p</= 0.0001, ****). Assumptions of equal variance are met. **F)** RT-PCR to check the quality of cDNA used for *doublesex* and *fruitless* assays, using primers for ribosomal protein S7. Three individual adult mosquitoes (3 biological replicates) were used for each genotype. Positive control (PC) and negative control (NC) templates were *Ae. aegypti* genomic DNA and H_2_O, respectively.

**Fig. S7.**
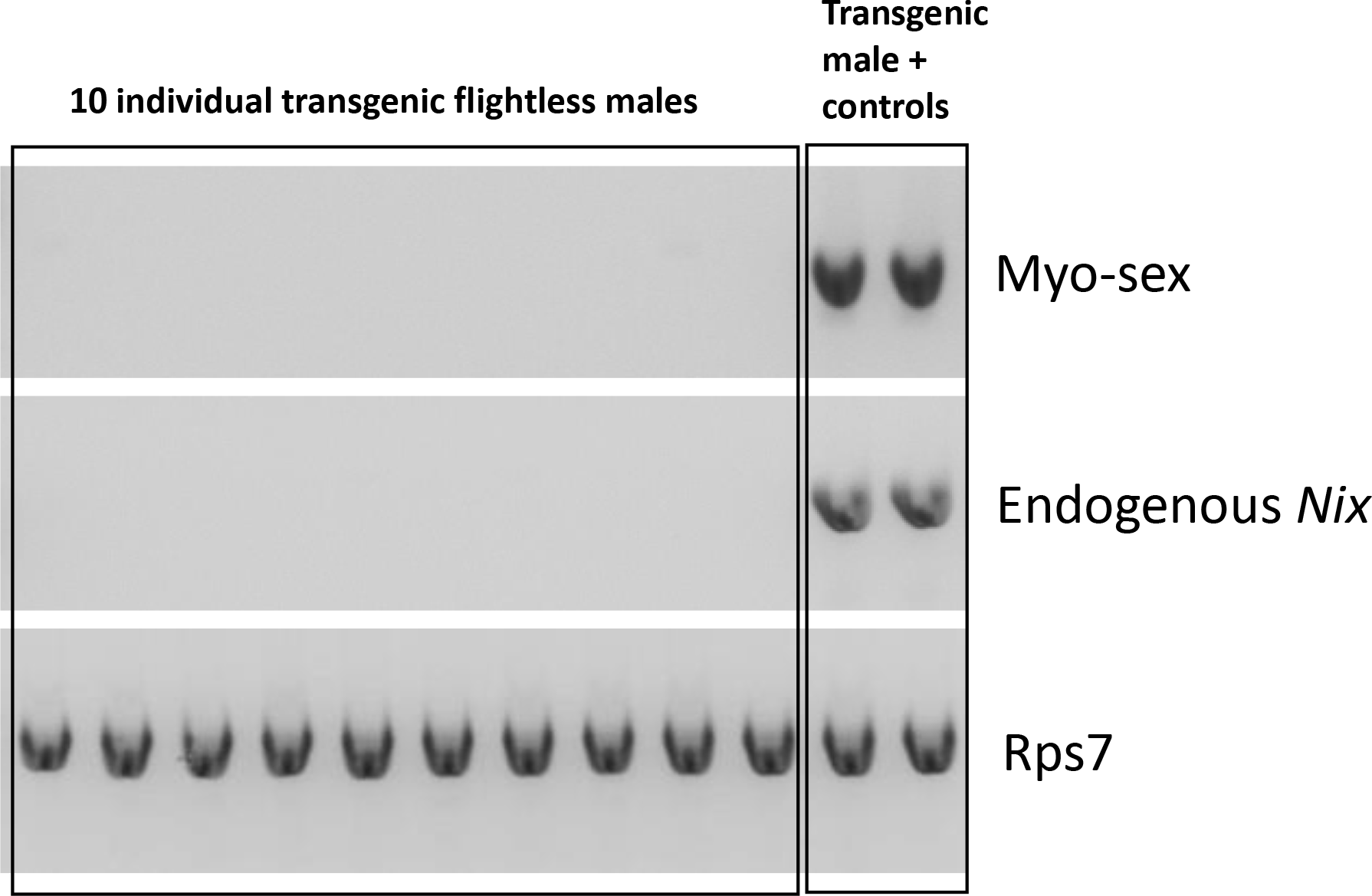
PCR assays used to determine that the flightless males in the N2 line are genetic females (m/m) that lack the M locus. These flightless males are progeny derived from forced mating. The absence of the M locus, as indicated by the absence of two M locus genes *myo-sex* and the endogenous *Nix*, is used to indicate the m/m genotype. Rps7 is used as the positive control. Size markers are not shown.

